# Effect of diet modification in intestinal *Escherichia coli* population and antimicrobial resistance

**DOI:** 10.1101/2021.04.12.439536

**Authors:** Fernanda Loayza-Villa, Daniela García, Alejandro Torres, Gabriel Trueba

**Affiliations:** Microbiology Institute, Colegio de Ciencias Biológicas y Ambientales, Universidad San Francisco de Quito. Diego de Robles y Pampite. Cumbayá- Quito. Ecuador; Carrera de medicina Veterinaria, Colegio de Ciencias de la Salud, Universidad San Francisco de Quito. Diego de Robles y Pampite. Cumbayá- Quito. Ecuador, E mail

## Abstract

The fluctuations in the number of some intestinal bacterial lineages may be associated with increased antimicrobial resistance and disease. Adaptation to a given environment may select bacterial mutants that have reduced ability to adapt to new environments and changes in diet have been associated with alterations in microbiome taxon composition. We wanted to see the effect of diet change in linage composition and antimicrobial resistance profiles of numerically dominant *E. coli*. We subjected 50 chickens from an industrial operation (under corn-based diet supplemented with antimicrobials) to 2 antimicrobial-free diets; one based on corn and the other based on alfalfa. Fecal samples were obtained from all animals at arrival and after five weeks under different diets. Five *E. coli* colonies (from each sample) were subjected to genetic typing and antimicrobial susceptibility testing. We observed high diversity and high turnover rate of numerically dominant *E. coli* strains from animals from both diet groups. We did not find differences in antimicrobial resistance profiles in isolates from different diet groups. Our results suggest that there is high diversity and high turnover rate of *E. coli* strains in the intestines regardless of the diet. Chicken intestines seemed to contain many *E. coli* lineages able to thrive in different substrates. The absence of differences in antimicrobial resistance among bacteria, from animals in different diets, may indicate that the carriage of antimicrobial resistance genes does not affect the bacterial ability to adapt to different substrates.

## 1. Introduction

A warm-blooded animal may harbor more than 10 different commensal *E. coli* lineages in the intestine, some of these lineages are numerically dominant (Lautenbach et al., 2008). The relative abundance of some *E. coli* lineages is critical because many strains carry virulence genes (Madigan, et al. 2015) and/or troublesome antimicrobial resistance (Wang, et al., 2020). The relative lineage abundance in the intestine, at any given time, may depend on their aptitude to use intestinal substrates, differential to bacteriophages, protozoan predation, and host immunity (Brito, et al., 2016; Sutton & Hill, 2019; Tenaillon et al., 2010; Wildschutte et al., 2004)

It has been shown that different diets have a profound impact on the relative abundance of intestinal bacterial species (Chung et al., 2016; Frese et al, 2015; Gagnon et al., 2007; Niu et al., 2015; Rowland et al., 2018), however, little is known about the impact of diet change in bacterial lineages belonging to the same bacterial species. Different diets may affect the growth of different bacterial strains in the intestine; different members of the same bacterial species may have lost or acquired different metabolic properties through mutation or horizontal gene transfer (Hehemann, et al., 2010; Brito et al., 2016; Leiby and Marx, 2014). These acquired properties may facilitate or hamper the ability of some strains to use some substrates present in the diet. Genome analysis and culture experiments show that different *E. coli* strains have a variable aptitude to use different substrates (Baumler et al., 2011; Monk et al., 2013; Bouvet, et al., 2017; Barrera, et al., 2019); even a single *E. coli* strain passaged thousands of times in culture media produce descendants with different growth rates in different substrates (Leiby and Marx, 2014). Adaptation to some substrates may reduce their ability to proliferate in other substrates (Buckling, et al. 2003; Leiby and Marx, 2014).

Similarly, antimicrobial resistance genes may affect the ability of some members of the microbiota to replicate in the intestine. Genes involved in antimicrobial resistance cause fitness costs are eventually ameliorated by compensatory mutations (Andersson and Hughes, 2010; Andersson and Hughes, 2011; MacLean, et al., 2010). However, the compensatory mutations could also have fitness costs and may limit the bacterial ability to diversify and adapt to new environments or substrates (Buckling, et al. 2003). One example of this phenomenon could be the reduction of antimicrobial-resistant bacteria in the intestine after diet changes (Wu et al., 2016; Auffret et al., 2017; Liu, et al., 2019).

In this study, we aimed to observe how the *E. coli* lineages change in chicken intestines as diet is drastically altered. We analyzed the effects of a diet change in the relative frequency of numerically dominant *E. coli* strains chickens. We also assessed the effect of diet change in the frequency of antimicrobial-resistant phenotypes in numerically dominant *E. coli*.

## 2. Methods

### 2.1 Animals

All protocols of experimental design were approved by the ethics and biosecurity committee of the Animals Ethics Committee of Universidad San Francisco de Quito before the study.

One hundred Cobb 1day old chickens (vaccinated against Marek Gumboro, New Castle, and Bronchitis) were donated from an industrial operation where animals are fed with a corn-based diet. All animals were kept in the same diet (without antimicrobials) for 2 weeks before splitting the chickens into different study groups. To rule out potential effects of the surrounding environment, identical experiments were carried out in two locations (1.- USFQ 2.- farm). A randomized design was conducted with 4 homogeneous groups with 25 chicken; groups were kept separated through the experiment. In the 2 locations, one group of animals was feed with the conventional diet (D1), and the other group was feed with an alternative formula based on dry alfalfa pellets (D2), neither group received antimicrobials (Supplementary materials Table S1). This regimen was maintained for the following 5 weeks. Water was available *at libitum*. Each chicken was an experimental subject which has identified with a mark painted in the plumage. Fecal samples were collected from ten chickens from each pen.

### 2.2 Samples and phenotypic analysis

Fecal samples were taken from 10 randomly selected chickens which were marked for future identification. Samples were obtained from the same chickens at week 2 and week 7. Each chicken was separated in a clean cardboard box until a fecal sample was obtained in a sterile container and maintained in ice for transportation to the lab within one hour after collection. Samples were plated on MacConkey agar and incubated at 37°C for 18hours. Five lactose positive colonies were selected form the plate and β-glucuronidase activity was confirmed using Chromocult Agar (Merck).

### 2.3 Antimicrobial susceptibility test

Five confirmed *Escherichia coli* isolates from each plate were isolated and stored at - 80°C in TSB with 30% glycerol (Cho et al., 2007). Antimicrobial susceptibility tests were performed using AMP ampicillin (10mg), TET tetracycline (30mg), SXT trimethoprim-sulfamethoxazole (1.25/23.75mg), GEN gentamycin (10mg), AMC amoxicillin-clavulanic ac. (20/10mg), CIP ciprofloxacin (5mg), CHLOR chloramphenicol (30mg), IMP Imipenem (5mg), CF cefazolin (30mg), CAZ ceftazidime (30mg), FEP cefepime (30mg), and CTX cefotaxime (30mg) as representatives of the most used families of antibacterial drugs in health care (Eisenberg et al., 2012). The Kirby Bauer test was carried out following CLSI (Clinical & Laboratory Standards Institute) guidelines using clinical settings for sensible or resistant phenotype interpretation.

### 2.4 Strain Genotyping

To determine whether the alfalfa diet could change the numerically dominant *E. coli* lineages, we analyzed the nucleotide sequences of the *fumC* gene in all isolates as published previously (Barrera, et al., 2019), and some strains showing identical sequence were submitted to full multilocus sequence typing (MLST) analysis. Briefly, the DNA from each isolate was released by the boiling method (Dashti, et al., 2009), two colonies (from each isolate) were placed in a test tube with 1 mL of molecular grade water, paced in a heat block at 100°C for 10 min., transferred to an ice bath for 30 seconds, and centrifuged for five minutes at 1,680 x *g* and supernatants were stored at −20°C for further analysis. The *fumC* gene was amplified as described previously (Wirth et al., 2006). Potential clonal isolates carrying *fumC4* and *fumC11* were subjected to MLST to confirm clonality (alleles *adk, fumC, gyrB,icd, mdh, purA, recA*) as previously reported (Wirth et al., 2006). Amplicons were purified and sequenced using commercial service based on the Illumina MiSeq platform at Macrogen Inc. (Seoul, Korea). Allele and MLST were obtained using http://mlst.warwick.ac.uk/mlst/dbs/Ecoli) to define clonal relatedness (Wirth et al., 2006).

### 2.5 Statistical analysis

Overall antimicrobial resistance was estimated at the isolate level with 95% confidence intervals. The AMR prevalence from feed treatment subgroup-collection was also estimated at the isolate level. For multidrug-resistant (MDR) estimation, isolates with resistant phenotype for three or more antimicrobial families were assigned as MDR. Prevalence estimates were carried out using the SPSS software version 24.0. The antimicrobial resistance profiles between different time points (2 weeks and 7 weeks) or feed source (corn-based vs. alfalfa-based) were compared by using the χ2 test; p values <0.05 were considered significant. For reduction analytic dimensions, principal components analysis for categoric variables were applied using CATPCA version 2.0 by Leiden SPSS Group (Leiden University, The Netherlands). Data were analyzed in a scatter plotter using XLSTAT 2019.4.2 (Addinsoft). Sequence types were assigned to *E. coli* isolates with a probable clonal relation and phylogenetic analysis was conducted on Mr. Bayes Vs 3.2 bases on MCMC algorithms. Evolutionary diversity analyses were conducted using the Maximum Composite Likelihood model in MEGA X.

## 3. Results

We found a large diversity and high turnover rates of numerically dominant *E. coli* lineages; we found that the 243 isolates had a limited number of *fumC* alleles (n=47); *fumC11* (n=72) and *fumC4* (n=30) were the most common (Figure 1). We run MLST analysis in a subset of the most common alleles: *fumC11* (n=16) or *fumC4* (n=8). Most of the isolates sharing the same *fumC* allele belonged to different sequence types except for 5 isolates which were ST48 (*fumC4*) found in 4 different animals with corn diet at week 2 and 1 isolate, from a different animal in alfalfa diet, at week 7 (Supplementary materials Table S2). Principal components analysis did not show any association between MLST profile and diet (Figure 2). Phylogenetic analysis of concatenated MLST sequences failed to show any cluster preferentially associated with any diet (Figure S1). Shannon diversity analysis of *fumC* alleles in both populations showed an H value of 2.04 for isolates from a corn-based diet and 2.43 for isolates from an alfalfa-based diet.

**Figure 1.**
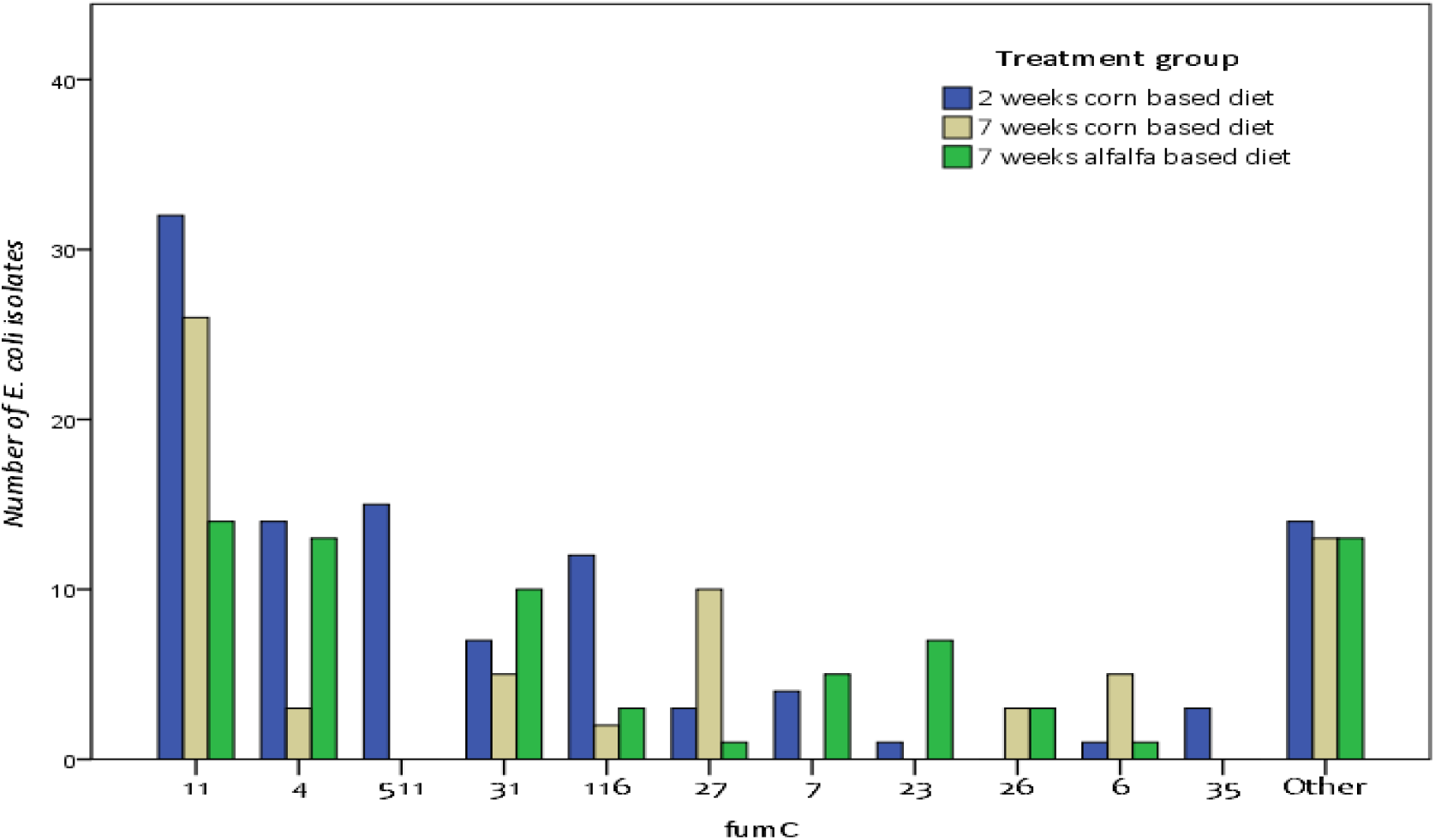
The number of *Escherichia coli* isolates carrying different *fumC* alleles from study groups. *E. coli* isolates (n=243) that were isolated from chicken feces; 106 from 2 weeks old chickens (blue bars); from 67 with corn-based feed (yellow bars) and 70 from chickens with alfalfa-based feed (green bars). Alleles with less than 1% of frequency where merge in “Other” category.

**Figure 2:**
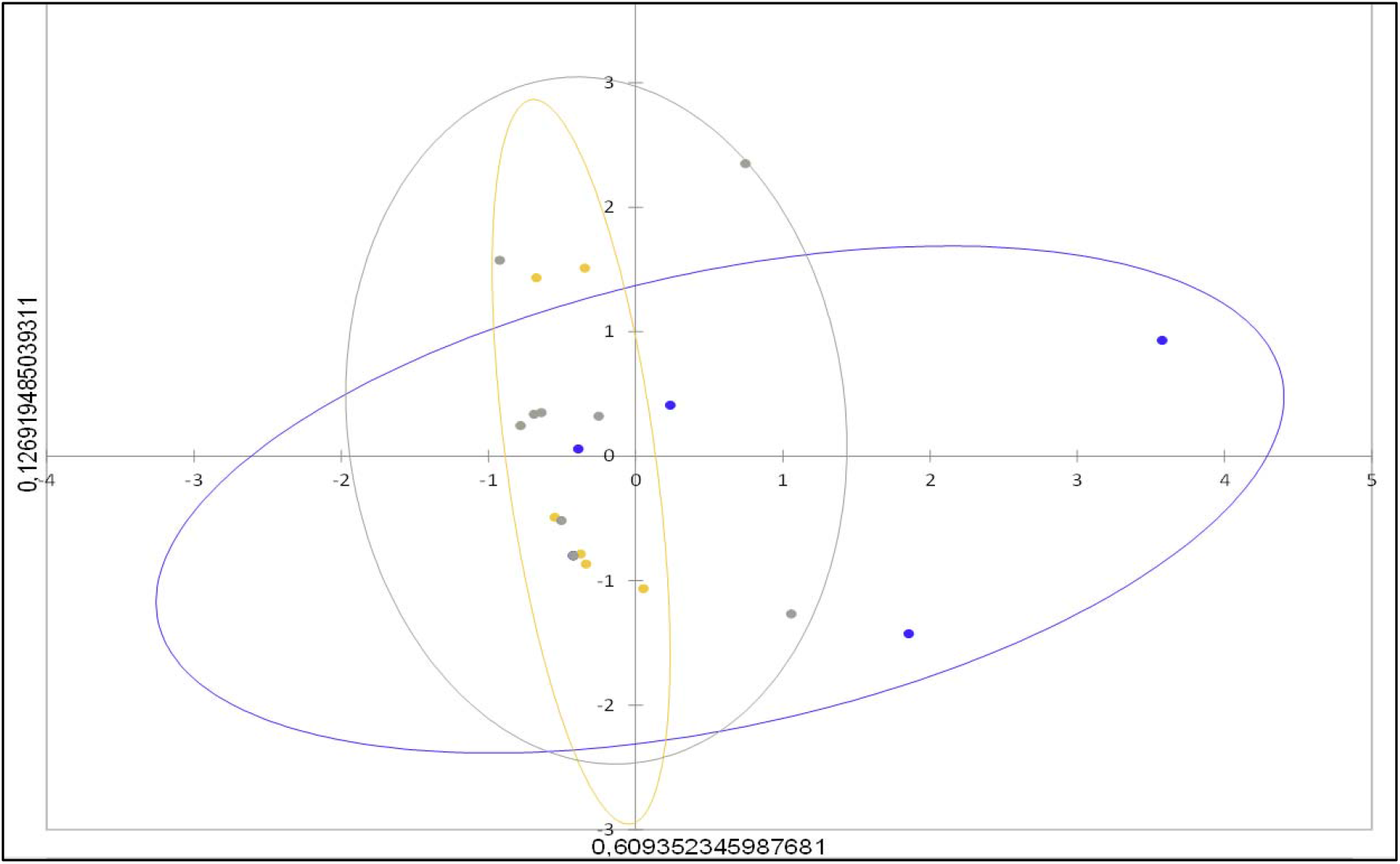
Principal components of 7 housekeeping genes profile of selected *Escherichia coli* isolates. 1. Blue dots represent isolates from 2 weeks chicken (basal), 2. Yellow dots represent isolated from 7-week chickens feed with a cornbased diet. 3. Grey dot represents isolates from 7-week chickens feed with an alfalfabased diet. Confidence intervals 95% based on chi-square are graph in colored ellipses according to the origin of isolates.

We did not observe differences in antimicrobial resistance phenotype in chickens in different diets (Supplementary materials Table S3) as 11. 1% of the strains from chickens with a corn-based diet and 18.7% of chickens in the alfalfa-based diet, were sensitive to all antimicrobials. After 5 weeks of intervention, 32 *E coli* isolates from chickens with a conventional diet were multi-drug resistant (MDR) compared with 30 isolates from chickens with an alternative diet. The principal component analysis of antimicrobial resistance phenotypes did not show differences between diet groups (Figure 3). We observed significant differences between 2 week-chickens and 7 week-chickens, regardless of the diet (X2; p< 0,05) (Figure 3; Supplementary materials Figures S2 and S3, Table S1). The most important resistance phenotype was tetracycline (TET; 11.25%) followed by tetracycline and cotrimoxazole (TET SXT; 8,25%). Antimicrobial resistance profiles *E. coli* were grouped in phenotype patterns. Some patterns were present in less than 1% isolates and represented the 15,75% of total phenotype patterns described (Supplementary materials Figure S2). MDR (resistance for 3 or more antimicrobials from a different family) was detected in 52.2% of *E. coli* strains. The most frequent combination was resistance to tetracycline, cotrimoxazole, chloramphenicol, and ciprofloxacin followed by tetracycline, cotrimoxazole, and ciprofloxacin combination with a 7 and 6.3% respectively.

**Figure 3.**
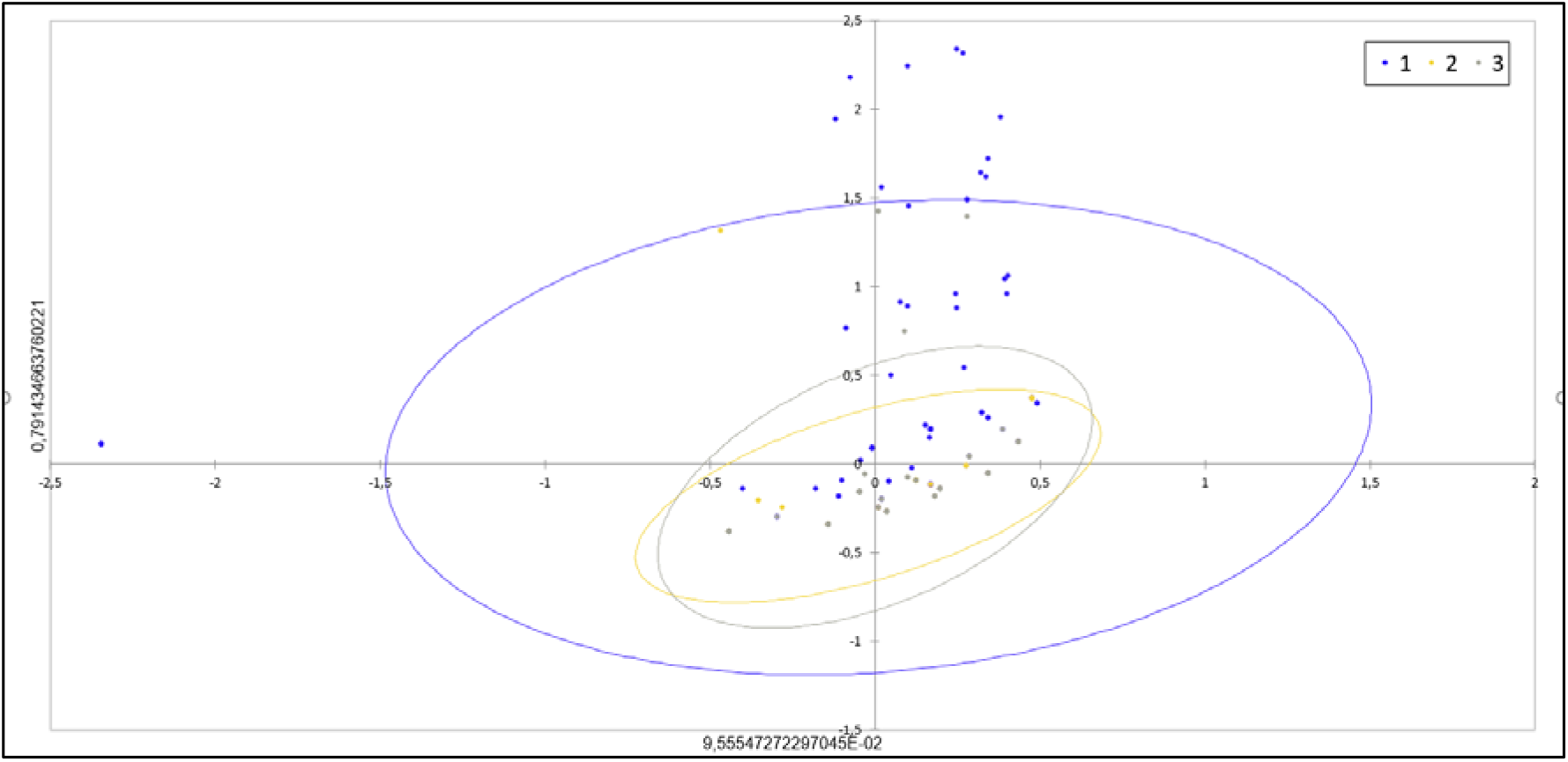
Principal components of phenotypic antimicrobial resistance profiles of 389 *Escherichia coli* isolates. 1. Blue dots represent isolates from 2 weeks chicken (basal), 2. Yellow dots represent isolated from 7-week chickens feed with a corn-based diet. 3. Grey dot represents isolates from 7-week chickens feed with an alfalfa-based diet. Confidence intervals 95% based on chi-square are graph in colored ellipses according to the origin of isolates.

## 4. Discussion

We showed high diversity and a high turnover rate of dominant *E. coli* lineages associated with chicken intestinal microbiota in the chickens in both diets (Figure 1). This finding agrees with previous reports showing high diversity and turnover rates of *E. coli* in human intestines (Richter, et al., 2018; Calderon, et al., 2020). We were not able to show any contribution of diet to strain diversity or turnover. We also failed to see clonal groups with a better aptitude to grow in alfalfa or corn (Supplementary materials Figure S1) which may indicate that genetically related strains may have different aptitude to grow in either of these substrates. The reason for this rapid turnover of dominant strains seems unclear but could be due to differential destruction of some *E. coli* lineages by bacteriophages (Sutton and Hill C., 2019), protozoa (Wildschutte et al., 2004), or host immunity (Tenaillion, 2010). It has been shown that many *E. coli* strains show the ability to adapt to different intestinal environments (generalists) as they move between different species of warm-blooded animals (Salinas et al., 2019; Moeller AH, 2018) and use diverse substrates.

We did not observe any significant variation in antimicrobial resistance in any group (Supplementary materials Figure S3), which may indicate that strains carrying antimicrobial resistance genes may not have any adaptive disadvantage when forced to grow in different substrates. Most antimicrobial resistance observed in these isolates (cotrimoxazole, chloramphenicol, and tetracycline) is widely disseminated in *E. coli* due to many years of antimicrobial pressure (Tedesse, 2012). Some antimicrobial resistance in *E. coli* emerged shortly after the invention of antimicrobials; sulfonamides were introduced in the 1930s and resistance was observed in 1950 (Tadesse et al., 2012); tetracycline was developed in 1948, resistance was observed in 1953 (Roberts., 1996); chloramphenicol was developed in 1947 (Tadesse et al., 2012) and resistance was found in 1955 (Watanabe, T., 1963). Plasmids carrying ARGs and bacterial chromosomes have evolved several compensatory mutations to ameliorate the fitness costs of ARGs (Andersson and Hughes, 2010; Andersson and Hughes, 2011). Contrastingly, antimicrobial resistance recently introduced in a bacterial species such as vancomycin resistance in *Enterococcus faecalis* and colistin resistance (mediated by the *mcr-1* gene) in *E. coli* have been easily reduced by eliminating the supplementation with these type of antimicrobials (Pantosti, et al., 1999; Wang, et al, 2020).

Finally, we observed a statistically significant reduction of isolates displaying antimicrobial resistance from 2 weeks to 7 weeks, in all experimental groups (Supplementary materials Figure S3.). We also observed differences in antimicrobial phenotype in strains collected from chickens at week 2 vs at 7 weeks(Figure 3). Previous studies have found that the proportion of antimicrobial-resistant strains is higher in 1-day- old than in older chickens (Hedman et al., 2019; Moreno et al., 2019). It has been proposed that antimicrobial-resistant *E. coli* lineages carried by 1-day chickens may gradually decrease as chickens grow (Moreno et al. 2019). This reduction may indicate that some plasmids and ARGs do cause fitness reduction as chickens grow.

The main limitation of this study is the low number of *E. coli* isolates analyzed from each chicken. Also, studying numerically dominant *E. coli* (Lautenbach, 2008) is an approximation to what happened to the E. coli population when the diet is changed, however, it is far less informative than a metagenomic analysis.

The persistence of antimicrobial resistance even in the absence of antimicrobials is a serious public health concern (MacLean, et al., 2010; Andersson and Hughes, 2011). Commensal *E. coli* plays an important role in the transmission of antimicrobial resistance from food-animals to humans (Berg et al., 2016; Hu et al., 2016), however very little is known about the dynamics of *E. coli* population in the intestines.

## Author contributions

**GT.** Principal researcher, edition of manuscript

**FL and DG.** All experiment performance, feeding animals, feces collection, microbiology analysis, statistics.

**AT.** Veterinary support and care of animals

## Supplementary Materials

### Tables

**Table S1.** Detailed analysis of diet formulation administered in each treatment groups

|  | Corn-based Diet | Alfalfa-based Diet |
| --- | --- | --- |
| Alfalfa pellets | 0% | 100% |
| Corn | 40% | 0% |
| Sorgo | 5% | 0% |
| Wheat | 12% | 0% |
| Rice | 3% | 0% |
| Soy | 5% | 0% |
| Soy paste | 24% | 0% |
| Fish floor | 3% | 0% |
| Bird floor | 3% | 0% |
| Salt | 0.376% | 0% |

**Table S2.-.** Sequence type from *Escherichia coli* strains. Alleles are described from each isolate. Treatment group: 1) 2-weeks-old chickens with a corn-based diet. 2) 7-weeks-old chicken with a corn-based diet and 3) 7-weeks-old chicken with an alfalfa-based diet. Also, phenotypic patterns of antimicrobial susceptibility from each isolate are described according to the antimicrobial discs used in Kirby Bauer test: AMP ampicillin (10mg), TET tetracycline (30mg), SXT trimethoprim-sulfamethoxazole (1.25/23.75mg), GEN gentamycin (10mg), AMC amoxicillin-clavulanic ac. (20/10mg), CIP ciprofloxacin (5mg), CHLOR chloramphenicol (30mg), IMP Imipenem (5mg), CF cefazolin (30mg), CAZ ceftazidime (30mg), FEP cefepime (30mg), and CTX cefotaxime (30mg).

| Treatment group | sample ID | Antimicrobial phenotype | Alleles |  |  |  |  |  |  | MLST |
| --- | --- | --- | --- | --- | --- | --- | --- | --- | --- | --- |
|  |  |  | adk | fumC | gyrB | icd | mdh | purA | recA | ST |
| 2 | 3.2 | AMC TET SXT CHLOR<br>AMP CIP | -196 | 11 | 55 | 101 | 113 | 40 | 38 | ni |
| 1 | 3.3 | TET SXT CHLOR AMP<br>CIP | 6 | 11 | 3 | 18 | 70 | 8 | 6 | ni |
| 1 | 8.2 | TET SXT CIP CHLOR | 6 | 11 | 4 | 8 | 8 | 8 | 2 | 48 |
| 2 | 8.3 | TET SXT CHLOR CIP | -199 | 11 | 4 | 8 | 8 | 1 | 4 | ni |
| 3 | 13.3 | TET CHLOR AMP | 6 | 4 | 4 | 10 | 7 | 8 | 14 | ni |
| 3 | 13.1 | AMC TET CTX FEP<br>SXT AMP | 6 | 11 | 14 | 16 | 24 | 8 | 6 | 2497 |
| 3 | 15.4 | TET IMP AMP | 6 | 11 | 4 | 8 | 8 | 8 | 2 | 48 |
| 1 | 14.1 | TET | 6 | 4 | 14 | 16 | 24 | 8 | 14 | 155 |
| 1 | 16.3 | AMC TET SXT AMP<br>CIP CF | 6 | 4 | 4 | 16 | 24 | 8 | 14 | 58 |
| 3 | 16.2 | TET SXT | 10 | 4 | 14 | 16 | 24 | 62 | 2 | ni |
| 1 | 18.3 | TET SXT CIP AMP GN | 6 | 11 | 4 | 8 | 8 | 8 | 2 | 48 |
| 3 | 18.2 | TET CIP | 10 | 4 | 4 | 8 | 8 | 8 | 4 | ni |
| 3 | 20.1 | TET AMP | 10 | 11 | 4 | 10 | 7 | 8 | 2 | 2705 |
| 1 | 20.3 | TET SXT AMP CIP | 6 | 4 | 14 | 1 | 20 | 62 | 7 | 345 |
| 2 | 21.4 | TET AMP | 6 | 11 | 5 | 8 | 8 | 8 | 2 | 6396 |
| 2 | 24.2 | TET SXT CIP | 10 | 4 | 4 | 10 | 24 | 8 | 14 | 5519 |
| 2 | 25.5 | TET CHLOR CIP | 24 | 11 | 4 | 8 | 8 | 8 | 73 | 73 |
| 2 | 26.4 | TET SXT CIP | 10 | 4 | 4 | 8 | 8 | 8 | 14 | 2883 |
| 2 | 26.5 | TET SXT CIP | 6 | 11 | 4 | 8 | 7 | 8 | 2 | 5224 |
| 1 | 27.4 | TET SXT CIP AMP GN<br>IMP CHLOR | 6 | 11 | 4 | 8 | 8 | 8 | 2 | 48 |
| 1 | 28.5 | AMC TET CTX CAZ<br>SXT AMP CIP CF | 10 | 11 | 14 | 8 | 8 | 8 | 313 | 4536 |
| 1 | 29.1 | TET SXT CIP | 6 | 11 | 4 | 8 | 8 | 8 | 2 | 48 |
| 2 | 30.3 | TET SXT CHLOR AMP | 6 | 11 | 14 | 10 | 7 | 8 | 2 | ni |
| 3 | 35.1 | TET SXT CIP | 24 | 11 | 4 | 8 | 8 | 8 | 2 | 43 |
| 3 | 39.1 | TET SXT CIP | 10 | 11 | 4 | 10 | 8 | 8 | 2 | 4704 |
| 3 | 40.4 | TET SXT | 10 | 11 | 4 | 8 | 8 | 1 | 2 | 1585 |

ni.- no identified

**Table S3.**
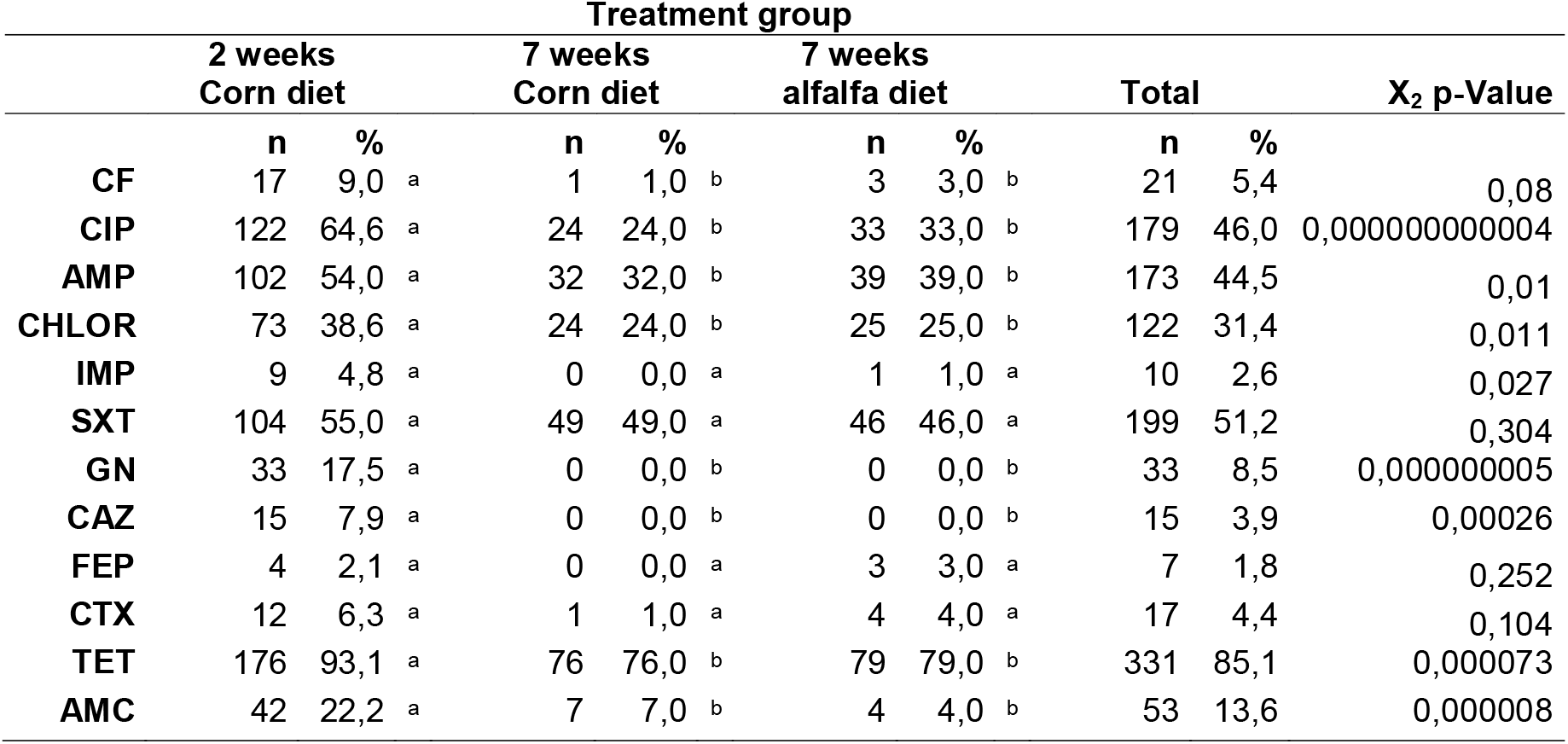
Frequency of resistance phenotype analyzed by the antimicrobial disc in Kirby Bauer test. In first column antimicrobial used are listed: AMP ampicillin (10mg), TET tetracycline (30mg), SXT trimethoprim-sulfamethoxazole (1.25/23.75mg), GEN gentamycin (10mg), AMC amoxicillin-clavulanic ac. (20/10mg), CIP ciprofloxacin (5mg), CHLOR chloramphenicol (30mg), IMP Imipenem (5mg), CF cefazolin (30mg), CAZ ceftazidime (30mg), FEP cefepime (30mg), and CTX cefotaxime (30mg). The letter after the percentage of resistant isolates in each group means the category according to X2 test. The same letter in three columns means that the proportions are not significantly different under the 0.05 level.

**Figure S1.**
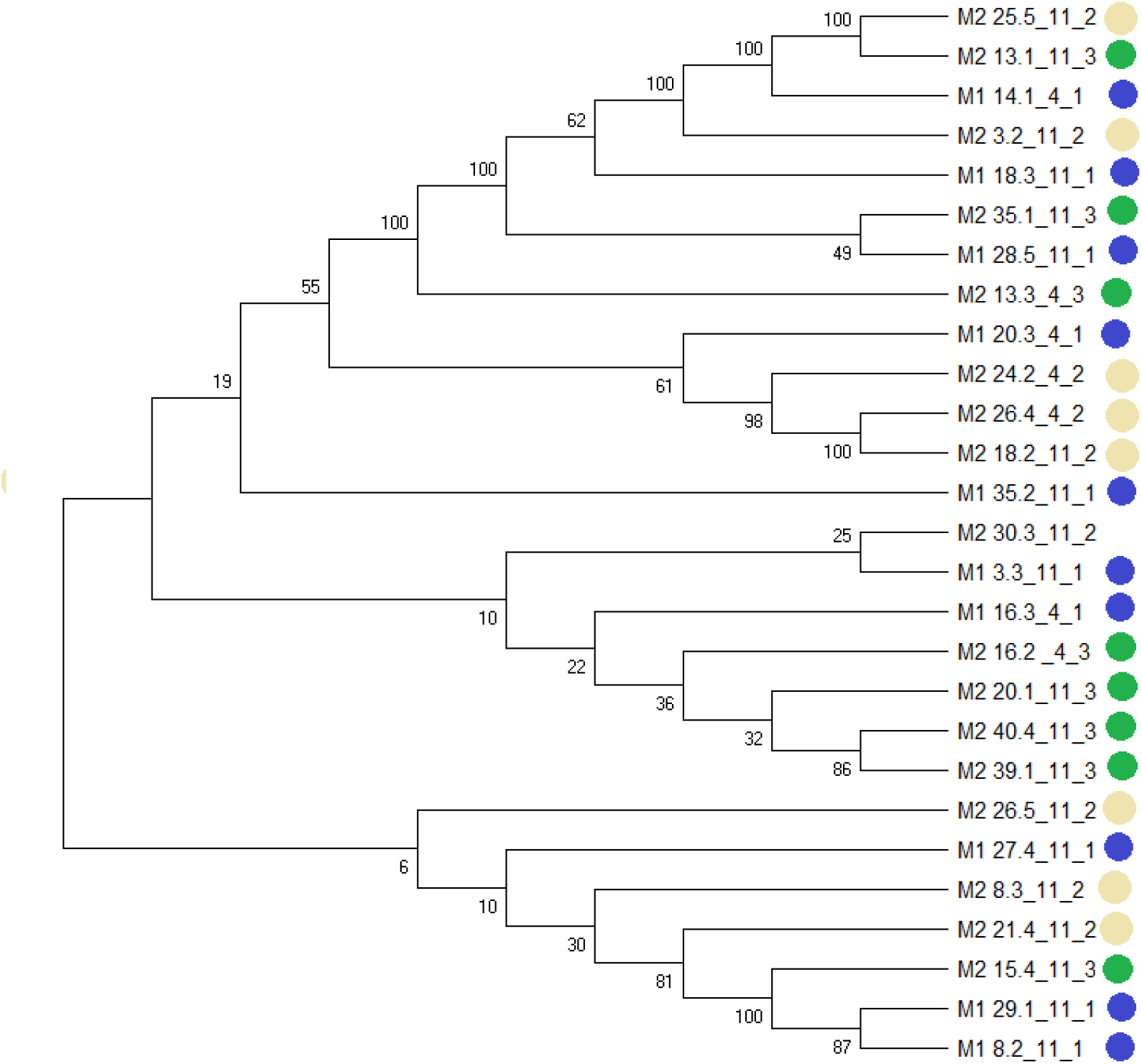
Bayesian phylogenetic tree was constructed from seven concatenated housekeeping genes used for MLST *E. coli* analysis. 1. Isolates from 2-week-old chicken (blue dots), 2. Isolates from 7-years-old chickens feed with a conventional corn-based formula (yellow dots) and 3. Isolates from 7-years-old chickens feed with an alfalfa-based formula (green dots). The identification label describes M1.-2 weeks old chicken, M2.- 7 weeks old chicken, followed by the ID number of each animal. After the underscore, *fum*C allele 4 or 11 are described. The last number after the second underscore confirms the treatment group.

**Figure S2.**
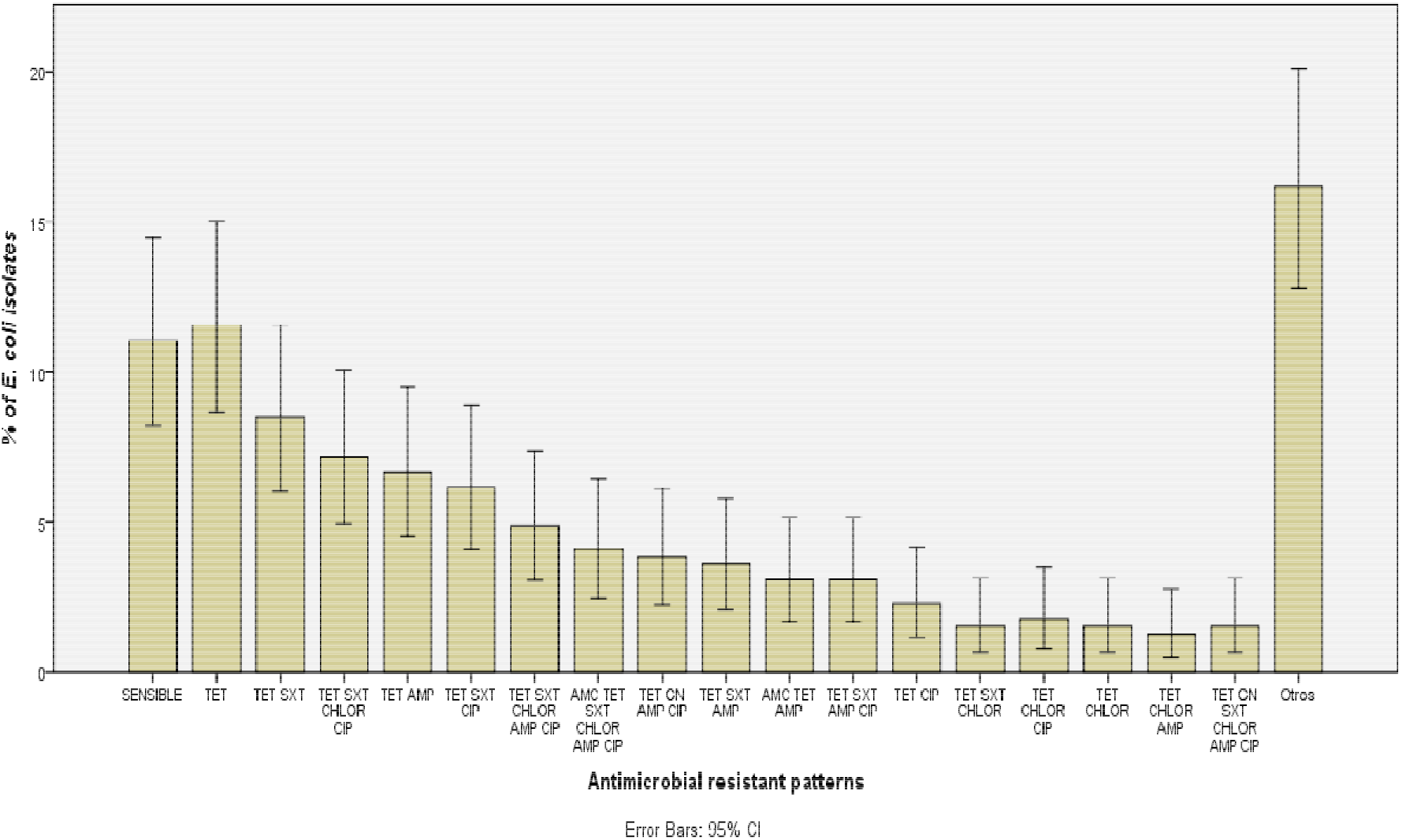
Overall frequency distribution of antimicrobial susceptibility patterns from 389 commensal *Escherichia coli* isolated from chicken feces in Mac Conkey Lactosa plates; 189 from 2 weeks old chickens, 100 from 7 weeks old chicken feed with a corn-based diet, and 100 7 week old chickens that were feed with alfalfa-based formula. Any diet has antimicrobial supplements. Kirby Bauer technique was performed for antimicrobial susceptibility testing (AST). AMP ampicillin (10mg), TET tetracycline (30mg), SXT trimethoprim-sulfamethoxazole (1.25/23.75mg), GEN gentamycin (10mg), AMC amoxicillin-clavulanic ac. (20/10mg), CIP ciprofloxacin (5mg), CHLOR chloramphenicol (30mg), IMP Imipenem (5mg), CF cefazolin (30mg), CAZ ceftazidime (30mg), FEP cefepime (30mg), and CTX cefotaxime (30mg).

**Figure S3.**
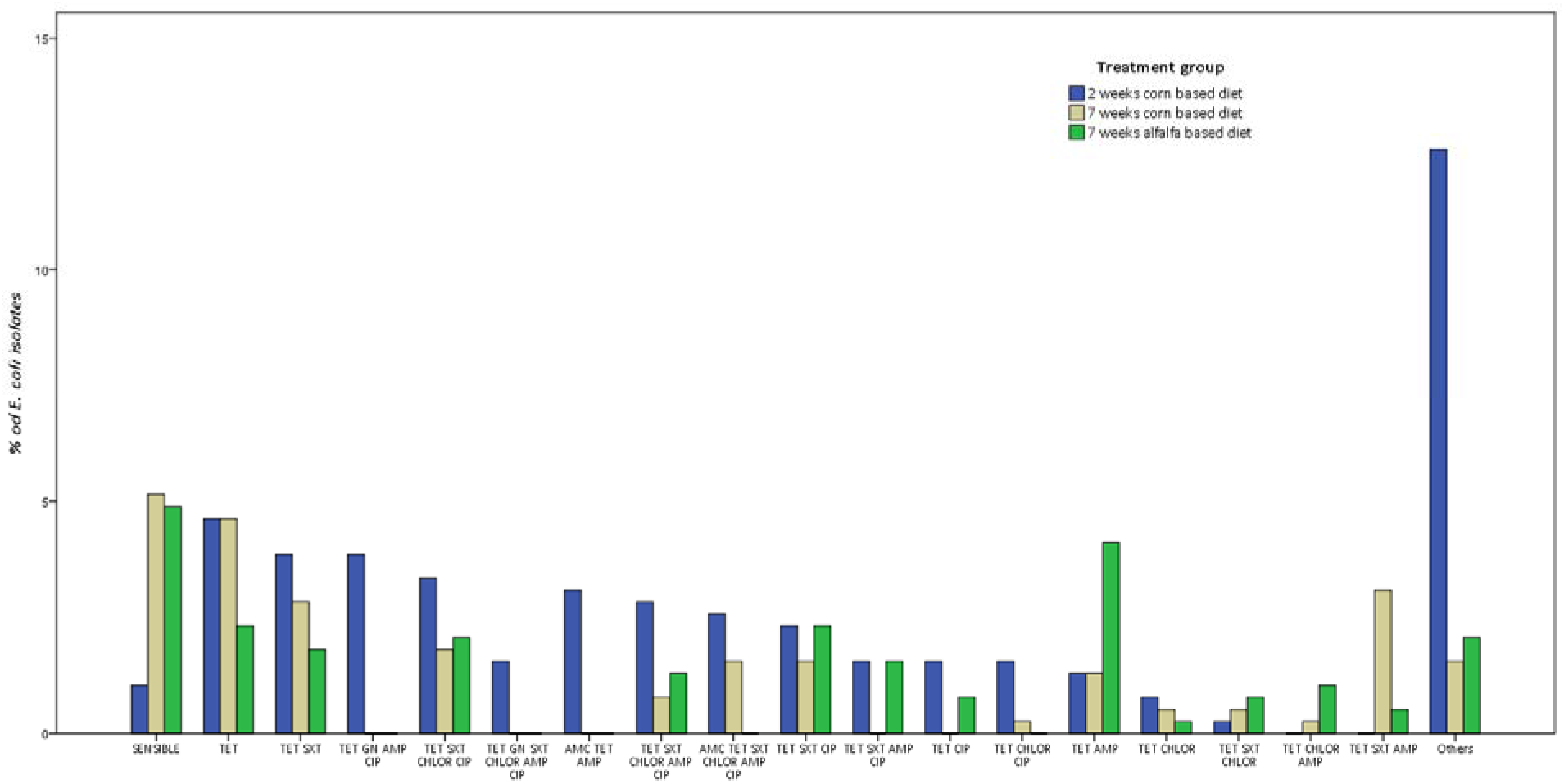
The number of *E. coli* isolates showing a specific phenotypic pattern of antimicrobial resistance. *E. coli* isolates (n=389) were isolated from chicken feces in Mac Conkey Lactosa plates; 189 from 2 weeks old chickens (blue bars) and 200 from 7 weeks old chickens, separated by feed administration 100 with corn-based feed (green bars), and 100 from chickens with *alfalfa-based* feed (yellow bars). Kirby Bauer technique was performed for antimicrobial susceptibility testing (AST) and antimicrobial-resistant phenotype patterns are shown. Patterns with less than 1% where merge in “Other” category. AMP ampicillin (10mg), TET tetracycline (30mg), SXT trimethoprim-sulfamethoxazole (1.25/23.75mg), GEN gentamycin (10mg), AMC amoxicillin-clavulanic ac. (20/10mg), CIP ciprofloxacin (5mg), CHLOR chloramphenicol (30mg), IMP Imipenem (5mg), CF cefazolin (30mg), CAZ ceftazidime (30mg), FEP cefepime (30mg), and CTX cefotaxime (30mg).

